# Arf6 is necessary for high level Wingless signalling during *Drosophila* wing development

**DOI:** 10.1101/2021.04.14.439835

**Authors:** Julien Marcetteau, Tamàs Matusek, Frédéric Luton, Pascal P. Thérond

## Abstract

Wnt signalling is a core pathway involved in a wide range of developmental processes throughout the metazoa. *In vitro* studies have suggested that the small GTP binding protein Arf6 regulates upstream steps of Wnt transduction, by promoting the phosphorylation of the Wnt co-receptor, LRP6, and the release of β-catenin from the adherens junctions. To assess the relevance of these previous findings *in vivo*, we analyse the consequence of the absence of Arf6 activity on *Drosophila* wing patterning, a developmental model of Wnt/Wingless signalling. We observed a dominant loss of wing margin bristles and Senseless expression in Arf6 mutant flies, phenotypes characteristic of a defect in high level Wingless signalling. In contrast to previous findings, we show that Arf6 is required downstream of Armadillo/β-catenin stabilisation in Wingless signal transduction. Our data suggest that Arf6 modulates the activity of a downstream nuclear regulator of Pangolin activity in order to control the induction of high level Wingless signalling. Our findings represent a novel regulatory role for Arf6 in Wingless signalling.

## Introduction

The ADP-ribosylation factor (Arf) family of small GTP-binding proteins is remarkably well conserved throughout the eukaryotes (Donaldson and Jackson, 2011). Arf6 is the most divergent of the Arfs, and localises to the plasma membrane and endosomes where it regulates various steps of endosomal trafficking and recycling (D’Souza-Schorey and Chavrier, 2006; Donaldson and Jackson, 2011). Previous *in vitro* studies have implicated Arf6 in the upstream stages of Wnt signalling (Grossmann et al., 2013; Kim et al., 2013; Pellon-Cardenas et al., 2013). However, a potential physiological, *in vivo*, role of Arf6 in Wnt signalling is yet to be addressed (Kim et al., 2013).

Despite the evolutionary distance between humans and *Drosophila*, Arf6 shares 97% sequence identity conservation between the two species **(figure S1A)**. Combined with the availability of powerful genetic tools, this makes *Drosophila* an ideal model in which to investigate the requirement for Arf6 in Wnt signalling in an *in vivo* context.

The *Drosophila* Wnt1 homologue, *wingless* (*wg*), is initially expressed throughout the wing primordium, and becomes progressively refined to a narrow stripe of cells of the presumptive wing margin late in larval development (Ng et al., 1996; Williams et al., 1993). The *Drosophila* wing has classically served as a developmental model of Wg signalling and has played a fundamental role in our understanding of Wnt/Wg signalling (Bejsovec, 2018; Jenny and Basler, 2014; Langton et al., 2016; Wiese et al., 2018). Canonical Wg signalling is contingent upon the stability of cytoplasmic Armadillo (Arm, the *Drosophila* β-catenin homologue) in signal receiving cells. In the absence of the Wg ligand, Arm is constitutively phosphorylated by the β-catenin destruction complex, consisting of the scaffold Axin, APC, and the kinases GSK3β and CK1 (Stamos and Weis, 2013), promoting Arm proteasomal degradation. The binding of Wg to the Frizzled 2 (Fz2) receptor and Arrow (Arr) co-receptor at the cell surface activates Dishevelled (Dsh), leading to the deactivation of the destruction complex and the stabilisation of cytoplasmic Arm (Swarup and Verheyen, 2012). Arm then translocates to the nucleus where it binds to Pangolin (Pan, a LEF/TCF homologue), converting it from a transcriptional repressor to an activator, and triggering the expression of Wg target genes (Mosimann et al., 2009; Schweizer et al., 2003).

High level Wg signalling is essential for the establishment and patterning of the wing margin (Couso et al., 1994; Jafar-Nejad et al., 2006; Phillips and Whittle, 1993). Cells flanking the wing margin respond to the local high levels of Wg protein by expressing the zinc finger transcription factor *senseless* (*sens*). Sens acts as the proneural factor for the anterior stout mechanosensory, and posterior non-innervated margin bristles (Jafar-Nejad et al., 2003; Jafar-Nejad et al., 2006; Nolo et al., 2000). Low level Wg signalling induces the expression of more sensitive target genes such as *distal-less* (*dll*) which is more broadly expressed in the wing blade (Neumann and Cohen, 1997; Zecca et al., 1996).

In this study we assessed the *in vivo*, developmental role of Arf6 in Wg signalling using a *Drosophila* model. *Arf6* mutants show a dominant loss of wing margin bristles and a concomitant loss of Wg-dependent Sens expression in the wing imaginal disc, phenotypes indicative of a defect in high level Wg signalling. In contrast to the previously suggested upstream roles of Arf6 in Wnt signalling (Grossmann et al., 2013; Kim et al., 2013; Pellon-Cardenas et al., 2013), our data indicate that Arf6 is necessary downstream of Arm stabilisation for the activation of high level Wg signalling. Moreover, we show that Arf6 acts upstream, or at the level of Pan activity. These findings represent a novel function for Arf6 in Wg signalling during wing margin development, and is the first demonstration of an *in vivo* role for Arf6 in Wg/Wnt signalling.

## Materials and Methods

### Fly genetics

Flies were raised in standard conditions. Crosses were carried out at 22ºC unless stated otherwise.

### Clone induction

Clones were generated by crossing males of either *FRT42B, Arf6*^*KO*^*/ CyO, Tb::RFP* or *FRT42B, Arf6*^*1*^*/ CyO, Tb::RFP* with virgins of *y, w, hsFLP; FRT42B, ubi-nlsGFP*. Heat shock induction was carried out for 30 minutes in a water bath at 37ºC, 48h after egg lay. Larvae carrying *Arf6*^*1*^ or *Arf6*^*KO*^ were selected based on the absence of *Tb*, then dissected and stained in wandering stage L3. Mutant clones were recognised based on the absence of a GFP signal.

### Fly stocks

The following fly stocks were used during this study: *w*^*1118*^ (Bloomington #3605) served as a wild-type control and the source of wild-type chromosomes. *Arf51F*^*GX16w*-^ (*Arf6*^*KO*^) (Bloomington #60585 (Huang et al., 2009)), *Arf6*^*1*^ (Dyer et al., 2007) (A kind gift from Marcos Gonzalez Gaitan) are both independently generated null alleles of *Arf6* lacking the full coding region. *Arf6*^*KO*^ was initially recessive lethal, so we introgressed both *Arf6* null alleles into a *w*^*-*^ background and reconfirmed the presence of the deletions by PCR. *Arf6*^*KO*^ and *Arf6*^*1*^ were maintained as a stock balanced over *CyO, Tb::RFP* (Bloomington #36336) to allow homozygous larvae to be recognised. High level Wg activation was induced using *UAS-dsh::myc* (Bloomington #9453), *UAS-sgg*^*A81T*^ (Bloomington #5360)(Bourouis, 2002), *UAS-Arm*^*S10*^ (encoding Arm lacking amino acids 37-84 in the N-terminus, Bloomington #4782) (Pai et al., 1997), *vgMQ-arm*^*NDel*^ (expresses a form of Arm lacking amino acids 1 to 138 from the N terminus, Bloomington #8370) or *UAS-axin-RNAi* (Bloomington #31705). Induction of Wg signalling downstream of Arm stabilisation was achieved using *UAS-pan*^*VP16*^*::HA* (generated in this study, see methods below).

Wg signalling inhibition was achieved with *UAS-dsh-RNAi* (KK330205, VDRC), *UAS-arr-RNAi* (GD6707 and GD6708, VDRC) or *wg*^*CX4*^ (Bloomington #2980). Wild-type *sens* was over-expressed with *UAS-sens* (Bloomington #42209). The following Gal4 drivers were used to drive expression in the wing: *nubbin-Gal4* (expressed throughout the wing pouch) (Azpiazu and Morata, 2000), *C96-Gal4* (expressed in a wide stripe overlapping the D/V boundary) (Bloomington #43343), *tub-gal80*^*TS*^; *hh-gal4/TM6b* in the posterior compartment. Mitotic clones were induced using *y,w,hsFLP; FRT42B, ubi-GFP*^*NLS*^ (derived from Bloomington #5826), and *Arf6*^*KO*^; *FRT42B/ CyO, Tb::RFP* or *Arf6*^*1*^; *FRT42B/ CyO, Tb::RFP* (derived from Bloomington stocks #1956 and #36336).

The following independently generated EMS-induced *pav* alleles were used: *pav*^*B200*^ (Bloomington #4384)(Salzberg et al., 1994) and *pav*^*963*^ (Bloomington #23926)(Collins and Cohen, 2005).

### Generating pan^VP16^::HA

*pan*^*VP16*^*::HA* was generated in order to allow the induction of Wg signalling downstream of Arm stabilisation. The construct is conceptually based on a construct previously shown to act independently of enhanceosome components Legless (Lgs) and Pygopus (Pygo) (Thompson, 2004). A sequence encoding full length Pan, excluding the stop codon, followed by 3xHA flanked by GGGGS linkers, and finally the VP16 transcriptional activation domain was synthesised (GeneArt). The sequence was directionally subcloned into 5’ KpnI and 3’ XbaI into *pUAST attb L34* plasmid (Bischof et al., 2007). Purified maxipreps were injected into the *M{3xP3-RFP*.*attP’}ZH-68E* background (Bl# 24485) (Bischof et al., 2007) in order to generate third chromosome insertions.

### Antibodies

The following primary antibodies were used: rabbit anti-GFP (1:400, Life Technologies A6455), Guinea pig anti-Sens (1:1000, a kind gift from Hugo Bellen), rat anti-Distalless (1:100, a kind gift from Marc Bourouis), mouse Anti-Wg (1:100, DSHB 4D4), mouse anti-Arm (1:10 DSHB N2 7A1). Rat anti-DE-cadherin (1:50, DSHB DCAD2).

The following secondary antibodies were used: Goat anti-rabbit Alexa488 (1:500; Invitrogen A11034), goat anti-rabbit Alexa546 (1:500; Invitrogen A11035), donkey anti-mouse Alexa488 (1:500; InvitrogenA21202), donkey anti-mouse Alexa546 (1:500; Invitrogen A10036), donkey anti-rat Alexa488 (Invitrogen A21208), goat anti-rat Alexa546 (1:500; Invitrogen A11081) and TRITC-phalloidin (1:100; Sigma P1951-1MG)

### Wing imaginal disc preparation and imaging

Wandering stage L3 larvae were washed then dissected in ice-cold 1xPBS. Fixation was carried out for 20 minutes at room temperature in 3.7% formaldehyde with constant agitation. Samples were washed and permeabilised for 30 minutes in PBT (0.3% Triton X-100, 1x PBS) then blocked for 1h in blocking buffer (0.1% Triton X-100, 1% BSA, 1x PBS) at room temperature. Primary antibody incubations were carried out over-night at 4ºC in 200µl of antibody diluted in blocking buffer. Samples were washed 3x 20minutes in PBT, then incubated for 1 hour at room temperature with secondary antibodies. Samples were washed in PBT then mounted in VECTASHIELD mounting medium (Vector Laboratories).

Images were acquired with a Leica TCS upright SP5 confocal microscope using a 40x objective (HCX PLAN APO; Numerical aperture of 1.3). The leica LAS AF software package was used for image capture (v2.6.3.8173). Images were analysed using FIJI (Schindelin et al., 2012) and the data analysed and visualised in R (R Core Team, 2013).

### PCR validation of Arf6 deficiencies

Genomic DNA was extracted from individual flies. Flies were crushed in PCR tubes using a pipette tip containing 50μl of squashing buffer (10mM Tris-HCl, 1mM EDTA, 25mM NaCl and 200 μg/mL proteinase K). Samples were incubated at 37ºC for 30 minutes then heat inactivated at 95ºC for 2 minutes using a thermocycler. 1μl of the resulting extraction was used as the PCR template.

The deficiency described for *Arf6*^*1*^ was validated using PCR **(figure S1B’)** and the primer combinations shown in **(figure S1B)**. *Arf6*^*KO*^ has previously been characterised in (Huang et al., 2009). 2x GoTaq Green Master Mix (M7121, Promega) was used for the PCR reactions. The following primers were used to validate the *Arf6*^*1*^ allele:

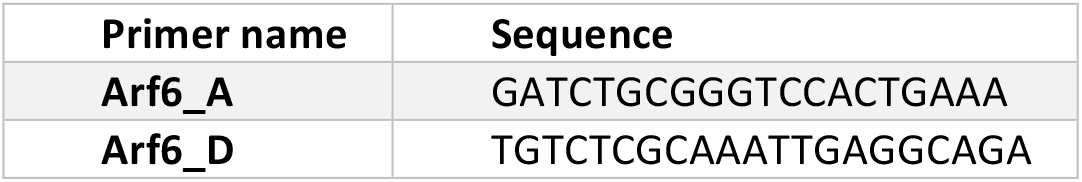

### Adult wing dissection

Adult flies were collected in ethanol at least 12h following emergence to ensure their wings had fully expanded and dried. Wings were removed at the hinge in ethanol, dried on blotting paper, then mounted in a drop of Euparal (Carl Roth #7356.1) and left to cure over-night on a slide heating plate set at 60ºC. Wings were imaged using a Leica DM2000 with an attached Leica DFC7000T camera. Wings were excluded from quantifications if damage to the wing margin prevented bristle quantification.

### Quantification and statistical analysis

The numbers of both ectopic and stout wing margin bristles **(figure S1C)** were quantified manually using the cell counter plugin in FIJI (Schindelin et al., 2012). Statistical analyses and plotting were carried out in R (version 3.6.3)(R Core Team, 2013). Stout bristle counts in the anterior margin were modelled using Generalised Linear Models (GLM) with Poisson errors. Counts of ectopic bristles were modelled using GLM with Poisson errors, and Zero-Inflated negative binomial GLMs were used to compensate for the large number of wings that contain no ectopic bristles (in **figures 3B and D**)(Zeileis et al., 2008). Pairwise contrasts between genotypes for both margin and ectopic bristles were calculated using the emmeans package (Lenth, 2021). The p values resulting from multiple comparisons were corrected for Type 1 error using Bonferonni correction. Plots were generated using the GGPLOT2 package and exported using the egg package (Auguie, 2019; Wickham, 2009). Sample sizes are marked on the plots or provided in figure legends.

## Results and Discussion

### Arf6 is necessary for wing margin patterning

We observed a dominant reduction in the number of bristles throughout the wing margins of adult flies heterozygous for the amorphic *Arf6* alleles, *Arf6*^*1*^ and *Arf6*^*KO*^ (Dyer et al., 2007; Huang et al., 2009) (**figure 1A, A’’**) (see **figure S1C** for wing margin bristle patterning). This phenotype was strongly enhanced in homozygous *Arf6* mutants (**figure 1A, A’’)**. The trans-allelic combination of *Arf6*^*1*^ and *Arf6*^*KO*^ resulted in a comparable phenotype to the respective homozygotes **(figure 1A, A’’)**, showing that the loss of the DNA region common to both deficiencies is responsible for the phenotype **(figure 1A, figure S1B)**.

**Fig. 1.**
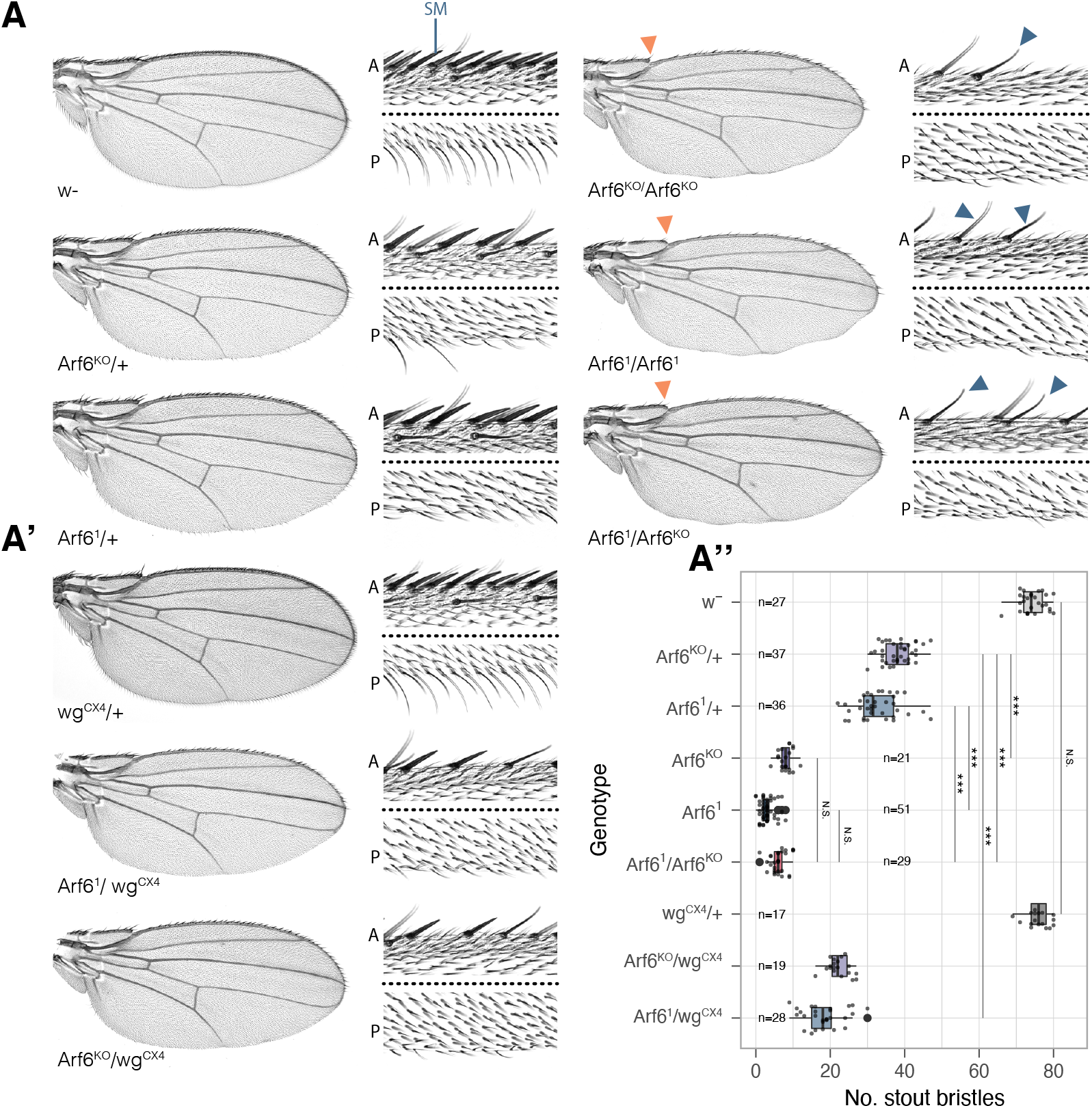
Dominant loss of wing margin bristles in Arf6 mutants. (**A-A’**) Representative wing blades and wing margins of control (*w*^*-*^), *Arf6*^*KO*^, *Arf6*^*1*^ and *wg*^*CX4*^ mutants and their genetic interactions. Zooms of the anterior (A) and posterior (P) wing margins are separated by a dashed black line. Slender chemosensory bristles are still present in the homozygous Arf6 mutants (solid blue arrows) while stout mechanosensory bristles (SM) are almost all absent. The solid orange arrows indicate the loss of distal costa bristles in *Arf6* mutants. The number of SM is quantified in (**A’’**). SM counts were modelled using a GLM with Poisson errors. Significance values resulting from post-hoc pairwise contrasts are reported by the following abbreviations: N.S. = p > 0.05, * = p ≤ 0.05, ** = p ≤ 0.001, *** = p ≤ 0.001.

The patterning of the wing margin is coordinated by high level Wg signalling late in larval development (Couso et al., 1994; Jafar-Nejad et al., 2006). We therefore tested whether the *Arf6* mutant phenotype is sensitive to the level of Wg. Although the null *wg* allele, *wg*^*CX4*^, does not induce a dominant wing margin phenotype **(figure 1A’, A’’)**, when in combination with either heterozygous *Arf6*^*1*^ or *Arf6*^*KO*^, it strongly enhanced the *Arf6* wing margin phenotype **(figure 1A’, A’’)**. We did not observe notching of the wing margin, or morphological defects in the bristles in *Arf6* mutants (**figure 1A, A’)**.

### Wg-dependent Senseless expression is suppressed in an Arf6 mutant

The zinc finger transcription factor Sens acts as the proneural factor for the margin bristles, and is expressed in two narrow stripes flanking the wing margin in response to high level Wg signalling (Jafar-Nejad et al., 2006; Nolo et al., 2000) (**figure 2A**). Sens expression was strongly reduced throughout the wing margin in an *Arf6* mutant context, but not in the sensory organ precursor in which the expression of Sens is independent of Wg **(figure 2A’)**. The bristles induced by ectopically expressing Sens were not suppressed in the *Arf6* mutant, indicating that the loss of bristles was not due to a loss of Sens proneural activity **(figure S2A, A’)**.

**Fig. 2.**
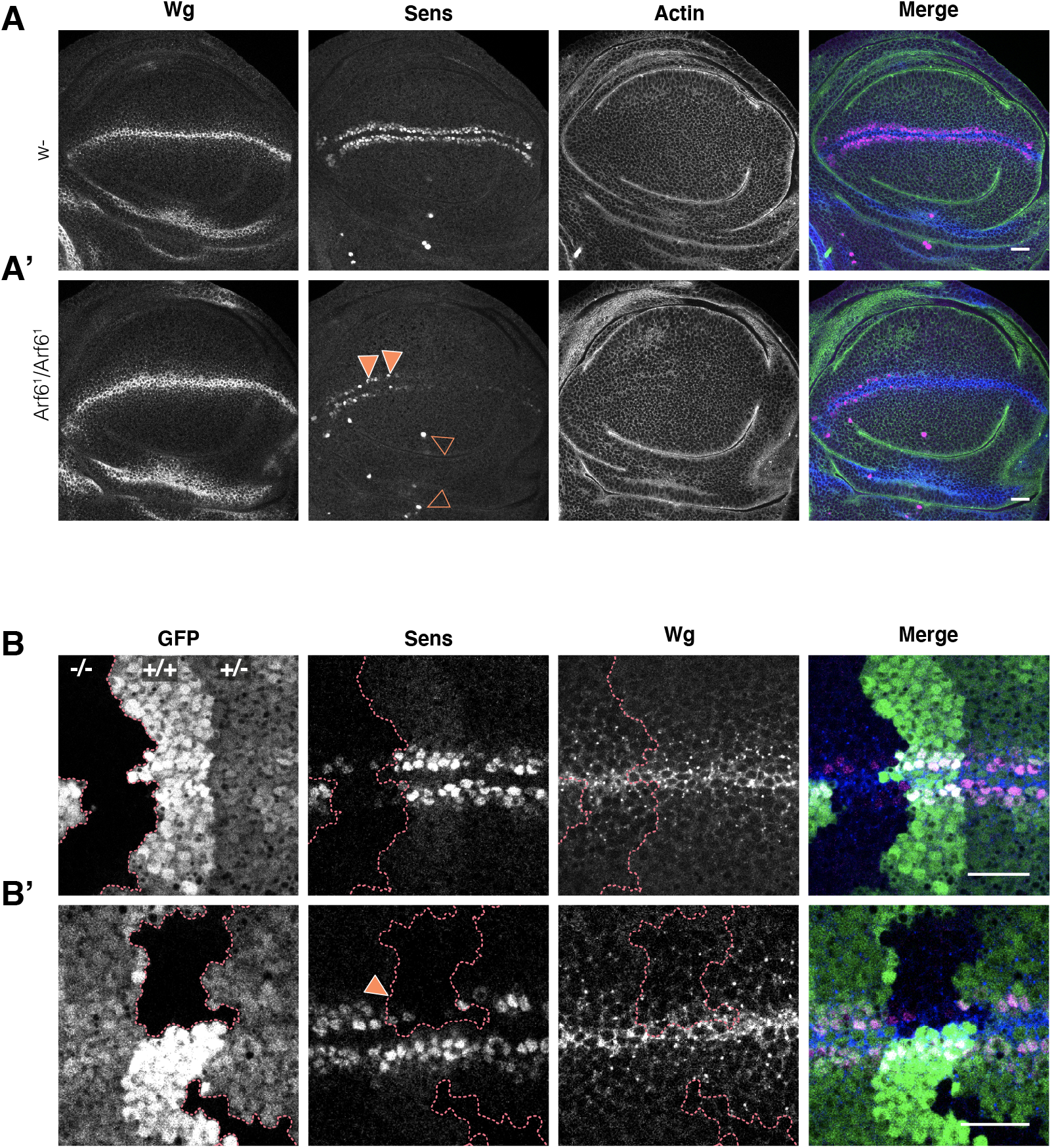
Level of Sens expression is strongly reduced in the absence of Arf6. (**A**) Wg and Sens staining in control (*w*^*-*^) and *Arf6*^*1*^ mutant. Anterior wing margin is to the left, posterior is to the right. (**A’**) Sens is almost completely absent in the posterior wing margin while Sens-positive cells are occasionally observed in the anterior wing margin (closed orange arrows) of the homozygous *Arf6*^*1*^ mutant. Sens is also observed in the prospective ventral radius and campaniform sensilla (open orange arrows). WT n=10, Arf6^1^ n=10. (**B**) Sens and Wg staining in *Arf6*^*1*^ mutant clones is marked by the absence of GFP (-/-). Heterozygous *Arf6*^*1*^ mutant and homozygous wild type tissue are marked by (+/-) and (+/+) respectively. In the merges, Sens is in magenta, Wg in blue, GFP in green (B-B’) and actin in green (A-A’). n=18. (**B’**) a strong reduction in Sens staining is observed in clones that do not enter the Wg expression domain. All scale-bars represent 20µm. n=19.

To test whether the loss of Sens expression is due to a defect in *wg* expression, we analysed the pattern of Wg in *Arf6* mutant wing discs **(figure 2A)**. The Wg stripe at the dorso-ventral (D/V) boundary was not disrupted by the loss of *Arf6*. Interestingly, the low-threshold Wg target Distal-less (Dll) was not reduced in *Arf6* mutant conditions **(figure S3A, A’, B)** indicating that Arf6 is not necessary for low level Wg signalling.

In order to assess whether Arf6 is required cell autonomously in Wg signal transduction, we generated random mitotic *Arf6*^*KO*^ clones which we then stained for Sens and Wg. Consistent with the dominant loss of bristles in *Arf6* mutants, we observed a strong reduction in Sens staining in homozygous *Arf6*^*KO*^ clones, an intermediate level in heterozygous tissue and unchanged level in the wild type tissue (**figure 2B, B’)**. Importantly, clones that overlapped with the *sens* expression domain, without entering the *wg* expressing margin cells, still induced a strong reduction in Sens expression (closed orange arrowheads, **figure 2B’)**, demonstrating that Arf6 acts cell autonomously in Wg receiving cells. Importantly, we did not observe ectopic Wg expression in *Arf6* clones near the wing margin (**figure 2B, 2B’**), nor wing notching in the *Arf6* mutant wing **(figure 1)** indicating that the integrity of the D/V boundary was not affected by the loss of *Arf6* (Rulifson and Blair, 1995; Rulifson et al., 1996). Altogether, these data show that while Arf6 is not required for the integrity of the D/V boundary, its activity is required cell autonomously for the transduction of high level Wg signalling controlling the expression of Sens necessary for wing margin bristle development.

### Arf6 is necessary downstream of Armadillo stabilisation

In order to determine the level at which Arf6 is required in Wg signal transduction, we began by activating the Wg signalling pathway in an *Arf6* mutant background. We suppressed the activity of the destruction complex by expressing a dominant-negative form of the *Drosophila* GSK3β homologue, *shaggy (sgg*^*A81T*^) (Bourouis, 2002) or knocking-down *axin*. Both treatments induce high level Wg signalling and the formation of ectopic bristles in the wing blade **(figure 3A, B)**. The number of ectopic bristles was dominantly suppressed in an *Arf6* mutant background **(figure 3A, A’, B, B’)**. These data indicate that the loss of bristles and Sens expression in the *Arf6* mutants is not a result of the hyperactivation of the Arm destruction complex, and suggest that Arf6 acts downstream of Arm stabilisation.

**Fig. 3.**
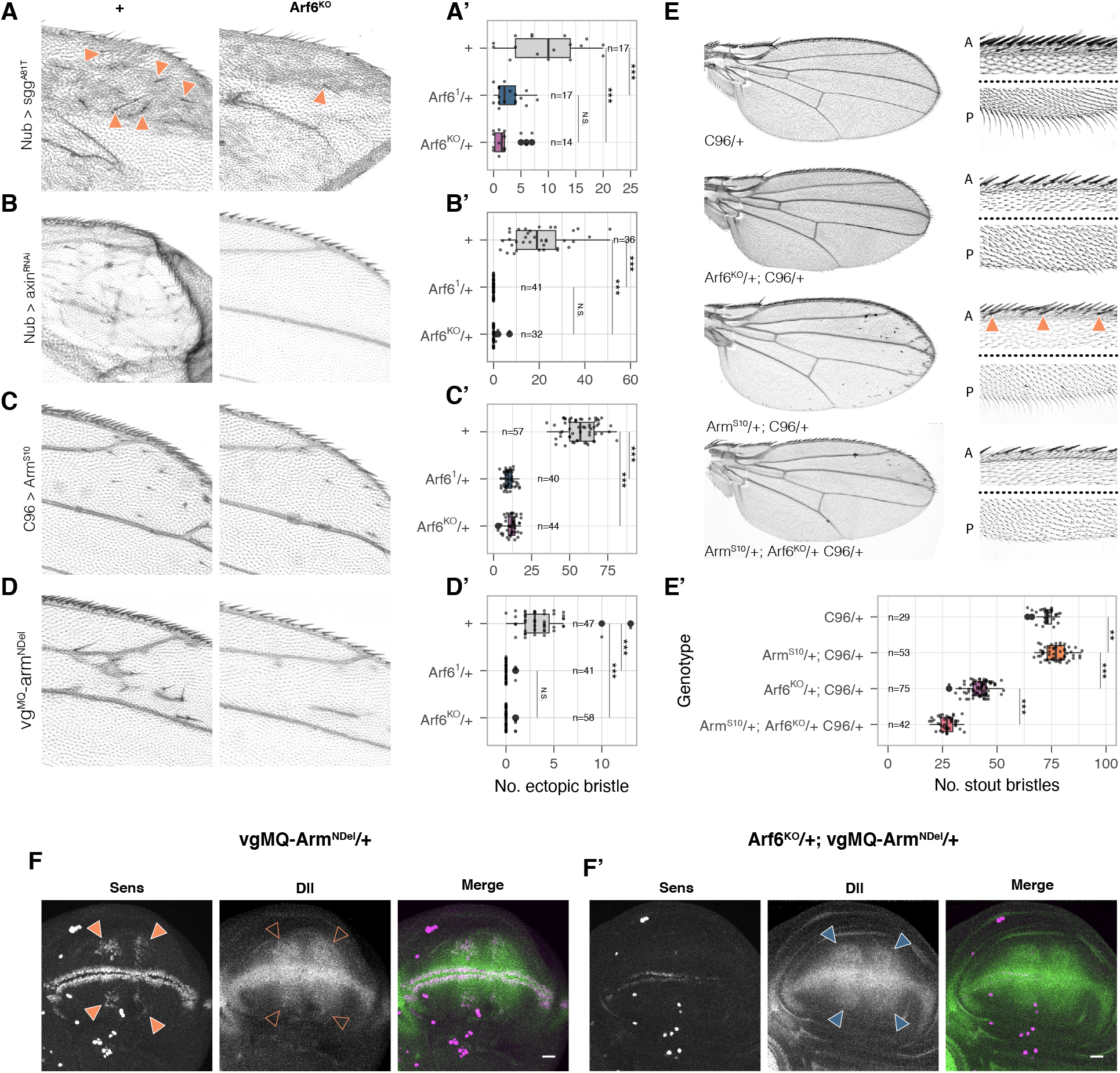
Epistatic analysis shows that Arf6 acts downstream of Arm stabilisation. (**A)** Dominant negative Sgg (*sgg*^*A81T*^) expressed with *nub-gal4* induces ectopic bristles (closed orange arrowheads), which are dominantly suppressed in the *Arf6* mutant background (quantification in **A’**). (**B)** Knock-down of *axin* induces ectopic bristles **(B’)** which are dominantly suppressed in the *Arf6* mutant background. (**C)** Arm^S10^ (expressed with *C96-gal*) and **(D)** *vgArm*^*NDel*^ (expressed under vestigial margin and quadrant enhancers) introduce ectopic bristles that are dominantly suppressed in the *Arf6* mutant background (quantified in **C’ and D’**). (**E)** Arm^S10^ expression with *C96-Gal4* at 25ºC enhances *Arf6*^*KO*^ margin phenotype, but introduces ectopic margin bristles in a wild type background (solid orange arrowheads). (**E’)** Quantification of stout mechanosensory bristles. Significance values resulting from post-hoc pairwise contrasts are reported by the following abbreviations: N.S. = p > 0.05, * = p ≤ 0.05, ** = p ≤ 0.001, *** = p ≤ 0.001. Ectopic bristles numbers were modelled using Zero Inflated negative binomial GLM. (**F)** *vgMQ-Arm*^*NDEL*^ induces ectopic Sens (closed orange arrowheads) and Dll (open orange arrowheads) in a wild type background. (**F’)** Ectopic Sens, but not Dll (closed blue arrows) is suppressed in the *Arf6*^*KO*^ background. In the merges, Sens is in magenta, Dll in green.

We next confirmed that Arf6 acts downstream of the stabilisation of Arm by expressing two constitutively active forms of Arm: Arm^S10^ and Arm^NDel^ (Pai et al., 1997). Importantly, these N-terminally truncated forms of Arm accumulate in the cytoplasm, triggering constitutive, high level Wg signalling in a ligand independent manner (Pai et al., 1997; Somorjai and Martinez-Arias, 2008). We expressed Arm^S10^ at the D/V boundary with the *C96-Gal4* driver, while Arm^NDel^ expression is directly driven by the vestigial quadrant and margin enhancers (subsequently referred to as *vgArm*^*NDel*^). Both Arm variants induced ectopic bristles in the wing blade **(figure 3C, C’, D, D’)**. Importantly, the bristles induced by *vgArm*^*NDel*^ were not dependent on endogenous Wg signalling (**figure S4A, A, B, B’, B’’)** and *vgArm*^*NDel*^ is active in canonical Wg signalling **(figure S4C)**. The ectopic bristles induced by both constructs were dominantly suppressed in the *Arf6* mutant background **(figure 3C, C’, D, D’)**. Moreover, Arm^NDel^ or Arm^S10^ did not rescue the wing margin bristles lost in the wing margin of *Arf6*^*KO*^ flies, and instead caused an enhancement of the *Arf6* mutant phenotype **(figure 3E, E’, figure S5A, A’)**. Over-expressing wild type *dsh* also induced ectopic bristles which were suppressed in *Arf6*^*KO*^ background (closed orange arrow, **figure S5B, B’)**. *dsh* over-expression also enhanced of the *Arf6*^*KO*^ phenotype (compare **figure S5C, S5B, S5C’)**. This is unlikely to be due to a dominant negative effect of Arm^S10^ or Dsh as expressing these constructs in a wild type background did not induce wing margin defects. Moreover, we did not observe a change in the levels of endogenous Arm and Cadherin at the adherens junctions in *Arf6* mutant clones (**figure S6A, A’**), suggesting that Arf6 does not regulate Wg signalling through the sequestration of Arm to the adherens junction in *Drosophila* (Grossmann et al., 2013; Pellon-Cardenas et al., 2013). Altogether, these data demonstrate that Arf6 is required downstream of Arm stabilisation in order to activate high level Wg signalling.

To test whether stabilised Arm had a generally reduced signalling activity in the *Arf6* mutants, we stained for both Sens and Dll in wing imaginal discs expressing *vgArm*^*NDel*^ in either a wild type **(figure 3F)** or heterozygous *Arf6*^*KO*^ background **(figure 3F’)**. Clusters of ectopic Sens positive nuclei were apparent far from the wing margin in wild type wings expressing *vgArm*^*NDel*^ (closed orange arrowheads **figure 3F)** accompanied by an upregulation of Dll (open orange arrowheads **figure 3F)**. Removing a single copy of *Arf6* led to an almost complete suppression of the ectopic Sens expression, including in the wing margin, but both the ectopic and endogenous Dll remained (closed blue arrowheads **figure 3F’)**. These data indicate that although Arm^NDel^ is still able to activate low level signalling in the *Arf6* mutant background, its ability to activate Sens expression is strongly attenuated. Importantly, the *Arf6* margin phenotype and suppression of high level Arm activity are independent of Arf1 **(figure S7A, A’, B, B’)** indicating that Arf1 and Arf6 play distinct roles in Wg signalling (Hemalatha et al., 2016).

Together, these results emphasise the specific requirement for Arf6 for the cell autonomous establishment of Sens expression in response to high level Wg signalling. The loss of margin bristles in the *Arf6* mutants is therefore likely to be due to a loss of the Sens-positive proneuronal clusters of the wing margin due to a lack of high level Wg signalling.

### Arf6 regulates Wg signalling at the level or upstream of Pangolin

The dominant suppression of N-terminally truncated Arm activity in *Arf6* mutants suggests that Arf6 could be involved in positively regulating canonical nuclear Wg signalling. Pavarotti (Pav), a MKLP1 homologue (Dyer et al., 2007; Makyio et al., 2012) has previously been shown to act in the nucleus as a negative regulator of Wg signalling during embryonic development (Jones et al., 2010). MKLP1 also recruits, and physically interact with Arf6 at the flemming body during cytokinesis (Makyio et al., 2012). We therefore hypothesised that Pav could provide the functional link between Arf6 and Wg signalling.

We began by testing whether the *Arf6* phenotype is sensitive to changes in the level of Pav. Pav is essential during cytokinesis (Adams et al., 1998), we therefore opted to use hypomorphic *pav* alleles (*pav*^*B200*^ and *pav*^*963*^) to avoid strong pleiotropic effects. Heterozygous *pav*^*B200*^ and *pav*^*963*^ flies in a heterozygous *Arf6* background provided a partial rescue of the number of wing margin bristles **(figure 4A, A’)** in the wing margin. These conditions did not induce cytokinesis defects or wing notching (**figure 4A, figure S8)**, consistent with Arf6 being dispensable for somatic cytokinesis in *Drosophila* (Dyer et al., 2007). The genetic interaction between *Arf6* and *pav* indicate that Arf6 could be regulating nuclear Wg signalling by modulating the non-canonical activity of Pav as a negative regulator of Pan activity (Jones et al., 2010).

**Fig. 4.**
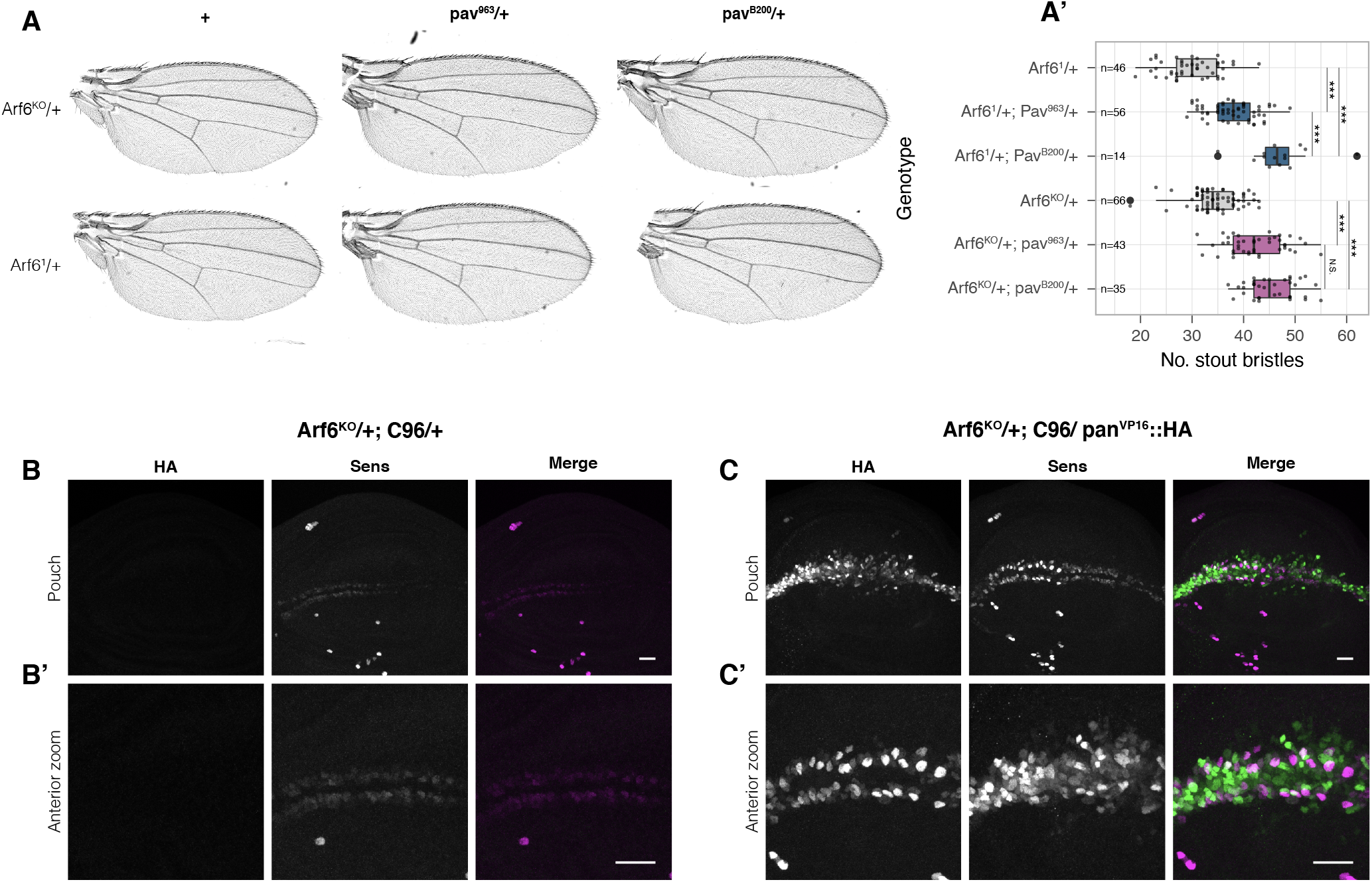
Pan-VP16 rescues Sens expression at the wing margin of Arf6 mutant. (**A**) The *Arf6* mutant phenotype is partially rescued in a hypomorphic *pav* background (stout mechanosensory bristles quantified in **A’**). (**B**) Wing imaginal discs showing Sens expression in *Arf6*^*KO*^*/+* and in (**C**) *Arf6*^*KO*^*/+* with *Pan-VP16::HA* expressed with *C96-gal4*. Anterior zooms of control and rescue discs are presented in **B’** and **C’** (in the merge, *Pan-VP16::HA* is magenta and Sens is green).

Once in the nucleus, Arm forms a complex with Pan, a TCF/LEF homologue forming the core of the enhanceosome (Gammons and Bienz, 2018). To determine whether Arf6 acts upstream of the enhanceosome, we generated a constitutively active form of Pan (Pan-VP16) **(figure S9A, S9A’)**. Expressing *pan-VP16* in a wild type background only induced low levels of ectopic Sens **(figure S9B**, closed orange arrowheads **S9B’)**, and was not sufficient to activate Sens expression far from the wing margin (open orange arrowheads, **figure S9B’)**, indicating that its activity still requires endogenous permissive signals. Expressing Arm^S10^ under the same conditions induced extensive ectopic Sens throughout the D/V boundary **(figure S9C, C’)**. Despite its greater ability to induce Sens expression, expressing Arm^S10^ with *C96-gal4* in a heterozygous *Arf6*^*KO*^ did not rescue Sens expression **(figure S9D, D’)**, whilst expressing *pan-VP16* in the same conditions resulted in a substantial rescue of Sens expression throughout the wing margin **(figure 4C, C’)**.

We have described a novel role for Arf6 in regulating high level Wg signalling, downstream of Arm stabilisation but upstream or at the level of Pan activity. The *Arf6* wing phenotype is particularly striking due to its dominance, as even mutations in core Wg pathway components do not dominantly induce wing margin defects (Couso et al., 1993; Couso et al., 1994). We therefore posit that the *Arf6* phenotype represents the derepression of negative regulators of the Wg pathway. Pav could provide a functional link between Arf6 and nuclear Wg signalling; Pav and Arf6 have previously been shown to cooperate during the spermatid cytokinesis (Dyer et al., 2007), however, Arf6 is dispensable for somatic cytokinesis (Dyer et al., 2007)(this study), and the suppression of Wg phenotype by Pav is independent of its role in cytokinesis (Jones et al., 2010). Arf6 could therefore conditionally sequester Pav in endosomal structures outside the nucleus, preventing its repressive activity in Wg signalling. Although previous studies have shown direct interactions between Arf6GTP to Pav/MKLP1 (Dyer et al., 2007; Makyio et al., 2012), further work is required to test whether this interaction is required for the loss of high level Wg signalling we observe in the *Arf6* mutants, and the mechanisms regulating Arf6 activity in this context.

## Acknowledgements

We thank all the members of the iBV ‘‘fly’’ community, Roland Le Borgne and Bruno Antonny for discussion. J.M. was supported by the French Government (National Research Agency, ANR) through the “Investments for the Future” programs LABEX SIGNALIFE ANR-11-LABX-0028 and IDEX UCAJedi ANR-15-IDEX-01 and by Fondation pour la Recherche Médicale.

## Author contributions

J. M. conducted all experiments except for Fig. S9 C-D performed by T. M. Experiments were design and discussed by J.M., T.M., F.L. and P.P.T. J.M. wrote the paper. All authors commented on the manuscript versions.

